# Quantitative multiplexed imaging of chromatin ultrastructure with Decode-PAINT

**DOI:** 10.1101/2022.08.01.502089

**Authors:** Hiroshi M. Sasaki, Jocelyn Y. Kishi, Chao-ting Wu, Brian J. Beliveau, Peng Yin

## Abstract

Super-resolution chromatin imaging techniques are emerging as powerful tools for spatial genomics. However, it remains challenging to visualize the genome on the order of tens of nanometers, a scale that can reveal rich structural detail about the nuclear organization. Here we introduce Decode-PAINT, a multiplexed in situ method, which leverages rapid diffraction-limited pre-decoding and DNA-PAINT (DNA points accumulation for imaging nanoscale topography). Decode-PAINT visualizes multiple discrete genomic regions at a resolution close to the nucleosome scale with a simple and easy-to-implement optical setup, reduces total imaging time by omitting multiple rounds of single-molecule recording, and enables the interrogation of the ultrastructure of the targets with <12 nm lateral and <20 nm axial resolution. We showcase the ability of our approach to perform multiplexed, high-resolution profiling of chromatin structure by performing targeted imaging of nine regions of active and inactive X chromosomes in individual cells. In short, Decode-PAINT accelerates in situ structural genomics.

## Introduction

Eukaryotic cells can package genomes with linear lengths of a few meters into a nucleus that is a mere ~10 μm in diameter. Despite this dramatic difference in length scales, the genome organization within the nucleus is non-random. Chromosomes occupy discrete nuclear space and adopt specific 3D conformations such as territories, compartments, and domains across the wide range of genomic scales^1–3^. The 3D organization of chromosomes regulates many nuclear processes such as transcription, duplication, and repair and plays crucial roles in our health and disease^4–6^.

Two major technologies have revolutionized our understanding of 3D genome organization: the chromosome conformation capture- (3C-) family of methods and single-cell imaging approaches such as fluorescent DNA in situ hybridization (FISH). 3C-family techniques, particularly one of the derivatives called Hi-C, rely on proximity ligation of digested genomic DNA and detect chromatin contacts at a length scale of megabases to a few hundred bases in a high-throughput manner. These allow us to uncover essential organizational elements such as loops, domains, and compartments in the genome^2,3,7^. However, despite considerable interest in recent years, many open questions remain about the 3D genome organization, particularly in individual cells, as 3C-family assays typically require populations of millions or more cells and single-cell Hi-C assays are still challenging due to their limited number of readouts. Imaging approaches such as DNA-FISH depend on the hybridization of oligonucleotide probes to specific genomic DNA sequences and have the potential to address this limitation, as they allow visualization of the spatial organization of specific genomic loci in single cells.

The utility of FISH techniques for interrogating chromatin structure and organization used to be hindered by the optics of light microscopy, as diffraction limits the maximum achievable resolution (~200 nm in XY and ~500 nm in Z), which is considerably larger than many nuclear structures. Breakthroughs in chromatin imaging in combination with Oligopaint FISH^8,9^ include highly-multiplexed sequential imaging^10–15^, direct single-molecule localization microscopy (SMLM)^11,16–18^, and a combination of both. In particular, sequential imaging methods allow us to map genomic loci by tiling them in short sections (2 kb–1 Mb) and use only the centroid positions of the imaged spots to reconstruct the chromatin paths in up to tens of thousands of cells with a relatively inexpensive cost^10–15,19–21^. Thus, these approaches are more suitable for addressing questions in chromatin traces, such as enhancer-promoter contacts and their dynamics during development and across cell types^12,13^. On the other hand, the SMLM methods locate the positions of single probes on the genomic targets using single-molecule photoswitching, enabling to visualize the 3D organization of the genome as point clouds beyond the aforementioned limitation. These approaches have addressed questions in chromatin packing organization, such as distinct compaction and intermixing for different types of epigenetic states and spatially segregated globular TAD-like structures observable in single cells^11,15–18^.

For most SMLM techniques, however, the achievable spatial resolution is still limited to ~30 nm^11,16–18^ by the nature of photoswitching in these super-resolution methods, as they must rely on intrinsic photophysical characteristics of the fluorescent dyes, which define the upper limit of the number of detected photons from each single-molecule event. Moreover, photobleaching of out-of-focus fluorescent dyes before detection, low efficiency of probe hybridization, and sequence constraints on probe design due to repetitive elements that occur abundantly in the genome can fundamentally limit the number of probes detectable per target region. These limitations, therefore, make it particularly challenging to detect small single-copy genomic loci in greater depth on the order of tens of nanometers. This scale could reveal rich structural information for quantitative analysis, such as the difference between active and inactive X chromosomes in female mammalian cells^22^, a pair of monoallelically-expressed gene loci^23,24^, or interphase and mitotic chromosomes^23^ <u>and thus allow the direct investigation of the effects of chromatin compaction state and ultrastructure on biological processes such as transcriptional regulation in the native context of single cells</u>.

DNA-PAINT is an SMLM technique to localize single-molecule targets by transient binding of fluorescently labeled oligonucleotides, called ‘imager strands,’ present in the imaging buffer to complementary strands, called ‘docking strands,’ on the target to be imaged^25,26^. DNA-PAINT has several advantages over other SMLM methods including decoupling of single-molecule events from dye photophysics, higher programmability of kinetic parameters, and better multiplexing. As new fluorescent strands continuously bind to targets, photobleaching does not reduce sensitivity. As single-molecule blinking signals are generated by DNA hybridization, DNA-PAINT enables the fine-tuning of the binding and unbinding rates by changing its sequence design or buffer condition such as its salt concentration, thus allowing the detectable photons from a single binding event to be maximized and the turn-over rate to be dynamically matched to the density of the target sites in the sample without significant signal overlap^25,26^. Finally, the programmability of DNA hybridization allows the design of many orthogonal DNA-PAINT imager-docking pairs for a sequential imaging approach^25,26^ (Exchange-PAINT) to visualize multiple targets in the same sample using only a single fluorescent dye^26^. In brief, different targets in the same sample are labeled with orthogonal docking strands. The targets are then imaged by sequentially applying corresponding imager strands. Together, these aspects have enabled DNA-PAINT to be used to perform sub-10-nm spatial resolution imaging^27,28^, quantification of the number of target molecules *in vitro* and *in situ*^29^, and up to three-color super-resolution chromatin imaging^16,18^.

Here we extend the applicability of DNA-PAINT to super-resolution quantitative chromatin imaging to a spatial resolution higher than that has been reached before. We first address the technical challenges of combining exchange-PAINT with chromatin imaging. We then introduce a combined approach, which we term ‘Decode-PAINT’, that integrates one-color standard DNA-PAINT with rapid pre-decoding by multiplexed SABER-FISH to perform super-resolution quantitative chromatin imaging of multiple discrete genomic loci in a relatively rapid manner. By separately applying the SABER imaging to rapidly identify genomic positions and then performing a single round of DNA-PAINT single-molecule imaging on all regions simultaneously, Decode-PAINT preserves the key advantages of both standard and super-resolution microscopy. Decode-PAINT enables us to accelerate image acquisition approximately three-fold by omitting multiple rounds of time-consuming singlemolecule recording. In addition, Decode-PAINT localizes the probes of the multiple targets in the same sample using just one DNA-PAINT imager, which results in generating a directly comparable dataset of super-resolved chromatin loci for further quantitative analysis at the single cell level. We demonstrate the utility of quantitative super-resolution imaging with Decode-PAINT by imaging the mammalian active and inactive X chromosomes, as the epigenetic differences between these chromosomes play an important role in cellular homeostasis and development^22^. We have found that inactive X chromosome has a similar internal compaction level with active X, supporting the idea that the chromosome organization of inactive X is distinct from that of the standard heterochromatic regions. Decode-PAINT provides researchers with a new tool for spatial functional genomics readily usable by biologists with standard lab equipment to perform a systems-level analysis of the geometric features and the internal organization of the targeted multiple genomic loci in individual cells.

## Results

### Evaluation of Exchange-PAINT in chromosome imaging

We first performed 8-color Exchange-PAINT in 3D to visualize a continuous stretch of chromosomes in defined increments **(SFig. 1)**. We previously designed a large set of orthogonal 42-nucleotide bridge sequences for barcoding^30^. Leveraging these bridge sequences, we designed primary Oligopaint oligos to label an 800-kb genomic target (spot) of human chromosome 8 in 100-kb increments **(SFig. 1a)**. Each probe has a spot-specific 42-nucleotide bridge sequence to accommodate the spot-specific docking sites **(SFig. 1b–d)**. While we successfully achieved the three-dimensional structure of the eight of 100-kb target loci at super-resolution via eight rounds of sequential DNA-PAINT imaging **(SFig. 1e,f)**, we found that it comes with two major drawbacks. First, because of the relatively slow imaging speed of DNA-PAINT, multiple rounds of singlemolecule recording for Exchange-PAINT often result in total acquisition times of more than one day for a single nucleus (**SFig. 1e**). Second, as each imager-docking pair has different binding kinetics, it is difficult to directly compare the localization data acquired using different imagers, making it less suitable for multiplexed quantitative analysis of genome organization.

### Principle of Decode-PAINT

To address the drawbacks of DNA-PAINT in chromatin imaging, we developed a strategy to simultaneously improve imaging throughput and streamline single-molecule localization. The recent development of the signal amplification by exchange reaction (SABER) technique enables rapid and multiplexed diffraction-limited imaging of DNA, RNA, and protein targets^30,31^. Here we introduce a combined approach, which we termed Decode-PAINT **(Fig. 1)**. By separately applying the SABER imaging to rapidly identify genomic positions and then performing a single round of DNA-PAINT single-molecule imaging on all regions simultaneously, we are able to preserve the key advantages of both standard and super-resolution microscopy.

**Figure 1.**
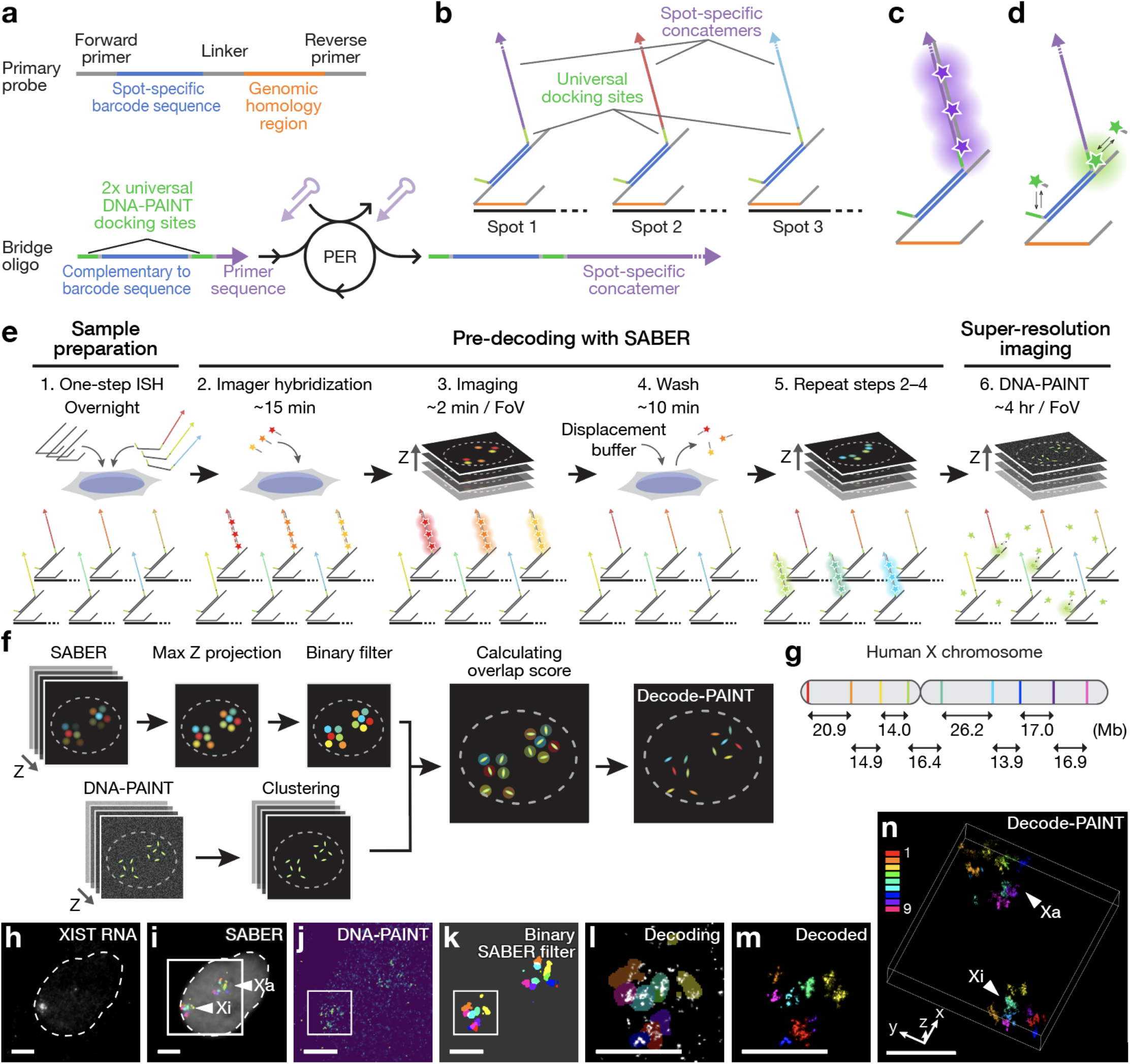
Decode-PAINT for visualization of discrete genomic loci at super-resolution. **a,** Primary probe and secondary bridge oligo designs. A primary probe has two main domains: spot-specific 42-nucleotide bridge sequence for barcode and genomic homology region for targeting. Forward and reverse primer sequences are used for probe preparation. A secondary bridge oligo carries a complementary sequence to a spot-specific barcode, two identical copies of a universal DNA-PAINT docking sequence, and a primer sequence at the 3’ end, which is extended by primer exchange reaction (PER)^52^. **b**, The tiling strategy for Decode-PAINT. While a unique concatemer sequence is assigned to each spot for SABER imaging, two copies of a universal DNA-PAINT docking sequence are shared by all genomic target spots, which reduces the number of long super-resolution imaging rounds required. **c,d,** Two types of readout from bifunctional bridge oligos. The bridge oligos can hybridize with fluorescently-labeled imagers for SABER (**c**) and generate singlemolecule signals via transient binding between DNA-PAINT imagers and docking sites (**d**). **e**, Decode-PAINT workflow. First, primary probe oligos and bridge oligos are hybridized to genomic targets in fixed samples following standard ISH protocols. The Z sections of the sample are then imaged by standard Exchange-SABER protocol to identify barcodes based on the fluorescence readouts (pre-decoding). *n* rounds of Exchange-SABER enable to identify 3*n* barcodes in an iterative manner. The sample is finally imaged with 1-color DNA-PAINT at super-resolution. **f**, Decoding workflow. The Z sections of SABER pre-decoded images are aligned using fiducial markers, converted to a maximum intensity projection of each channel, and then converted to a binary filter using Yen’s thresholding. DNA-PAINT data is segmented using the HDBSCAN clustering algorithm. The overlap score between the binary filter and the DNA-PAINT data is then calculated for each DNA-PAINT segment. **g**, Schematic of nine targeted regions along the human X chromosome. The genomic length of each target is 200 kb. **h**, XIST RNA FISH to identify Xi. **i–n**, Representative data of 9-color Decode-PAINT. Nine 42-nucleotide oligomer bridge sequences concatemerized with nine orthogonal PER primers were simultaneously hybridized. Scale bar, 4 μm.

In the Decode-PAINT scheme, we design bifunctional 42-nucleotide bridge oligos to label target-binding primary probes with a SABER concatemer strand and two copies of DNA-PAINT docking sequences **(Fig. 1a–d)**. Spot-specific barcodes are coupled with orthogonal SABER sequences to identify the barcodes of the targets from the readout of the SABER imaging. The experimental workflow can be divided into three steps **(Fig. 1 e; see Methods)**: (1) sample preparation, (2) pre-decoding, and (3) DNA-PAINT. First, both the primary probes and the bridge oligos are deployed onto the genomic targets in a single-hybridization incubation. Next, we perform multiplexed Exchange-SABER to identify each barcode using fluorescently-labeled SABER imagers prior to DNA-PAINT. Using three spectrally-resolvable dyes (ATTO 488, ATTO 565, and Alexa Fluor 647), *n* rounds of Exchange-SABER can identify *3n* barcodes in an iterative manner. Enhanced imaging with SABER enables rapid scanning of a field of view of 130 × 130 × 10 μm with three fluorescent channels at a 10-fold shorter camera exposure time (**SFig. 2a**), while achieving a comparable (ATTO 565 and Alexa Fluor 647) or better (ATTO 488) signal-to-noise ratio to non-amplified DNA exchange imaging (**SFig. 2b**). Moreover, ten minutes of washing with a formamide-containing buffer are sufficient to displace SABER imagers from the targets, which allows us to image tens of cells for diffraction-limited pre-decoding in a few hours (**SFig. 2a**). Finally, one-color DNA-PAINT localizes all the primary probes bound to the multiple targets at once.

Decoding is performed based on the overlap between the SABER signals and DNA-PAINT localization clusters using a custom Python pipeline **(Fig. 1f and SFig. 3a; see Methods)**. A pair of pre-decoding and DNA-PAINT data is registered using fiducial markers prior to the downstream processes. We generate a maximum intensity projection of each channel from the Z-stacks of the SABER images and then convert it to a binarized filter. After clustering of the DNA-PAINT localizations, we calculated the overlap score of each DNA-PAINT localization cluster with the SABER binarized filters to determine which targeted locus the cluster belongs to **(SFig. 3b; see Methods)**. Whereas the applicability of this method is limited to discrete genomic loci, it enables us to not only accelerate image acquisition approximately three-fold by omitting multiple rounds of time-consuming single-molecule recording (Supplementary Table 2) but also localize the probes of the multiple targets in the same sample using one DNA-PAINT imager, resulting in generating a directly comparable dataset of super-resolved chromatin loci in an improved throughput manner.

### Ultrastructural imaging of human X chromosomes

For proof of concept, we chose to target X chromosomes. In mammals, females carry two X chromosomes, while males inherit a single maternal X chromosome. For dosage compensation for the X-linked genes, one of the X chromosomes in females is transcriptionally silenced in the early developmental stage by way of the process called X-chromosome inactivation (XCI)^22^. Moreover, XCI leads to the condensation of the inactive X chromosome (Xi) into a cytologically detectable compact structure termed the Barr body, which is distinct from an elongated shape of the active X chromosome (Xa). Thus, Xi is thought to consist of facultative heterochromatin, resulting in the exclusion of transcription factors and a silenced state^22^. The major effectors of XCI include the XIST non-coding RNA, which is expressed on the only Xi and spreads in cis to coat the X chromosome resulting in the establishment and maintenance of XCI. Hi-C experiments performed in human and female mouse cells have revealed a global loss of domain structures and partition of Xi into two ‘super-domains’ of preferential self-interaction, which is separated by the DXZ4 macrosatellite^7,32–35^. On the other hand, a systematic single-cell analysis with diffraction-limited fluorescence *in situ* hybridization (FISH) has shown that differences of higher-order structures are visible at >10 Mb scale but are not reflected accordingly by enclosed, smaller segments^36^, which is in contrast to the prevailing view that Xi consists of facultative heterochromatin. To better understand the molecular architecture of Xi, it is important to further assess the geometric difference between Xa and Xi at super-resolution in individual nuclei, as this would allow a direct investigation of the packaging state of these chromosome to be investigated at the single cell level.

We designed a pool of oligo sequences targeting nine of 200-kb spots on the human X chromosome with spacer distances ranging from 13.9 to 26.2 Mb **(Fig. 1g)**. The library, which has been validated with FISH on mitotic chromosome spread^30^, allocates 1000 probes to each 200-kb spot (5 probe-binding sites per kb) for quantitative imaging. We first tested whether Decode-PAINT could be used to visualize multiple genomic targets in individual cells. Because chromosomes occupy limited territories in the nucleus, we expected signal spot decomposition to be particularly challenging for intrachromosomal genomic loci. We implemented the Decode-PAINT scheme by appending nine of 42-nucleotide bridge oligos coupled with orthogonal SABER barcode concatemers and the universal DNA-PAINT docking sequences shared by all genomic target spots **(Fig. 1a–d)**. We then performed simultaneous DNA-RNA FISH with two probe sets in XX 2N IMR-90 human fetal lung fibroblast cells, one of which targeted X chromosomal loci and the other XIST RNA transcripts as markers for Xi, allowing us to identify Xi prior to pre-decoding **(Fig. 1h)**. After iterative SABER imaging using wide-field epi-illumination for pre-decoding **(Fig. 1i)**, DNA-PAINT imaging was performed on the same microscope using oblique illumination covering ~3 μm from the coverslip surface with optical sectioning **(Fig. 1j)**. In total, nine colors (three hybridizations) of SABER were imaged in 4 hours, including the time taken for stripping, rehybridization, imaging >10 fields of view with z-stacks, followed by 4 hours of DNA-PAINT. Following the decoding workflow **(Fig. 1k–m)**, the targeted nine spots were successfully visualized **(Fig. 1n)**. Only 3% of localizations (614/19773) were found as unidentified due to overlapping signals in a diffraction-limited space.

### Quantitative and geometric analysis of active and inactive X chromosomes

Decode-PAINT can achieve high resolution and high-dimensional analysis of chromosomal nanostructures at the single-cell level. To illustrate this, we evaluated the performance of Decode-PAINT in the geometric comparison of chromosomal nanostructures between Xa and Xi (**Fig. 2, SFig. 4–6**). We targeted six of 200-kb spots on the X chromosome with spacer distances ranging from 16.4 to 40.3 Mb (**Fig. 2a**) and visualized them using Decode-PAINT from a total number of 95,000 frames of DNA-PAINT recording (**Fig. 2b-e; n_nucleus = 26 from 3 replicates)**. With our system, we achieved average XY and XZ localization precisions of 5.6 ± 2.5 and 7.8 ± 1.0 nm calculated by nearest neighbor-based analysis (NeNA)^27,37^, respectively, from singlemolecule localization events **(SFig. 4a, b)**, translating to ~11.5 and ~18.3 nm full width at half maximum (FWHM) supported resolutions. The spatial resolution of the Decode-PAINT data was evaluated as average 21.2 ± 4.8 and 28.8 ± 7.1 nm in XY and XZ, respectively, using a Fourier ring correlation (FRC) resolution analysis^38^ **(SFig. 4c, d)**. In addition, the FRC resolution analysis with downsampled data shows that resolution coarsens by only ~25% when between 70% and 100% data are used from the entire dataset (95000 frames), suggesting that the sampling efficiency is almost saturated **(SFig. 4e, f)**. Note that the FRC values are significantly lower than the supported resolutions calculated from NeNA due to the lack of periodicity in superresolved chromatin structures. The detection rate of each spot was ~60–90 % as some of the spots were located out of the Z range covered by oblique illumination and optical sectioning (**SFig. 5**).

**Figure 2.**
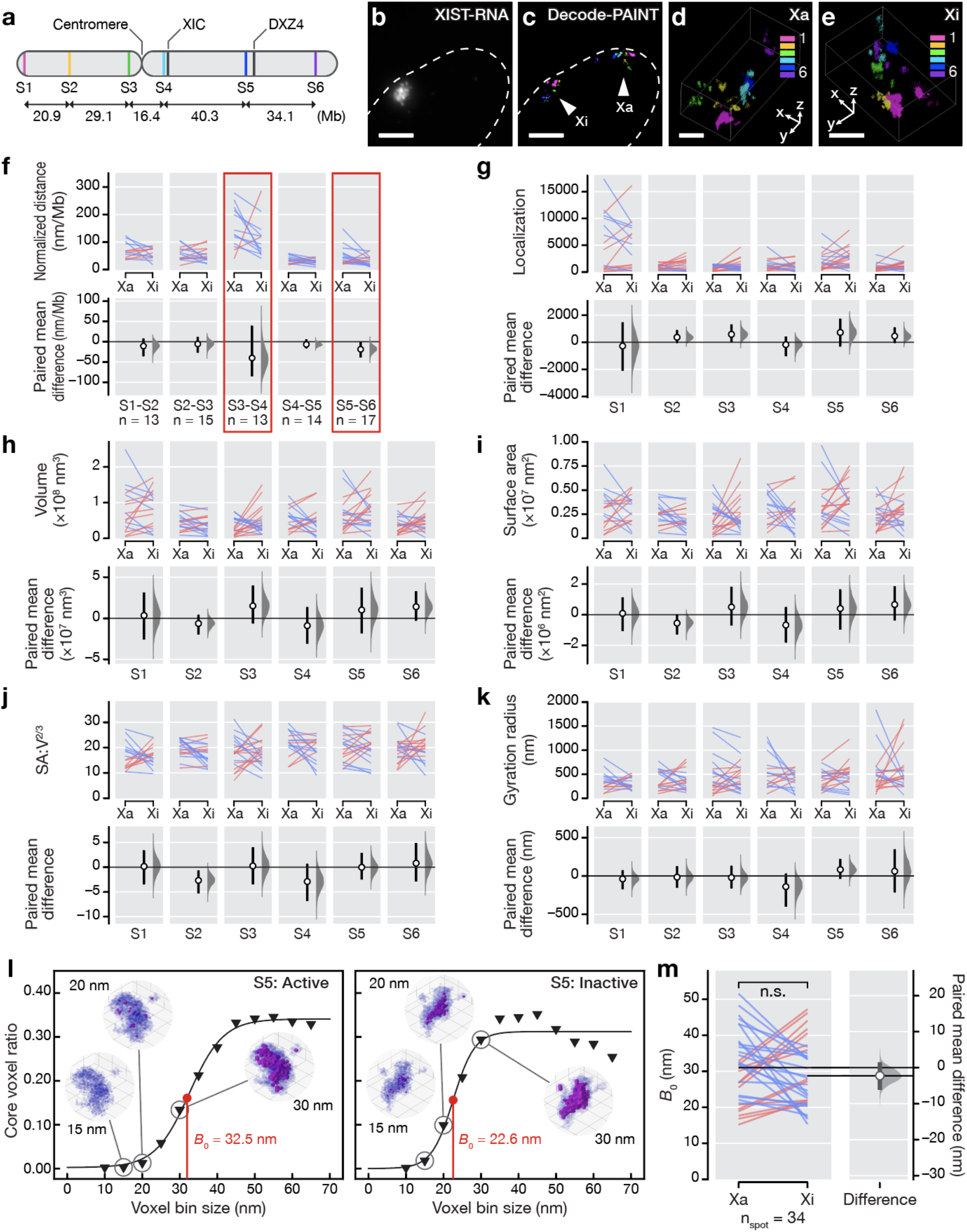
Geometric comparison between active and inactive X chromosomes with Decode-PAINT. **a**, Schematic of six targeted regions along the human X chromosome. The genomic length of each target is 200 kb. **b–e**, Representative data of 6-color Decode-PAINT with XIST RNA FISH. Scale bar, 4 μm. **f**, Paired estimation plots (slopegraphs and mean differences) of the normalized 3D distances between adjacent spot pairs at the single-cell pair level. Each pair of the slope represents the data points obtained from the active and inactive X chromosomes in the same cell. The physical 3D distances between the centroids of adjacent spots were divided by genomic distance. Red rectangles, *p* < 0.05 (the Wilcoxon signed-rank test). Each pair represents the data points obtained from the active and inactive X chromosomes in the same cell. **g–k**, Geometric analyses of active and inactive X chromosomes in single-cell pairs. Paired estimation plots of geometric features of active and inactive X chromosomes at the single-cell pair level. Each pair of the slope represents the data points obtained from the active and inactive X chromosomes in the same cell. Paired estimation plots of the following parameters were shown: the total number of localization recorded (**g**), volume (**h**), surface area (**i**), dimensionless surface-area-to-volume ratio (SA:V^2/3^) (**j**), and Gyration radius (**k**). **l**, Representative data of core voxel ratio plots from a voxel-based analysis. The left and right plots were generated from data of S5 of active or inactive X chromosome imaged in the same nucleus. Solid line, fitting curve with a sigmoid function. *B*_0_, the voxel bin size which gives the half maximum of the fitting curve, representing an apparent compaction level of the imaged spot. **m**, A paired estimation plot of *B*_0_. Each slope represents Xa-Xi paired data obtained from the same nucleus. All slopegraphs and paired mean difference plots were generated using DABEST^50,53^.

In our analysis, we first measured the physical 3D distances between the centroids of adjacent spots and divided them by genomic distance to obtain normalized distances (**Fig. 2f**). Significant distance differences (p < 0.05, the Wilcoxon signed-rank test) were found only between segments S3–S4 and S5–S6, which encompass the centromeric region and the DXZ4 locus, respectively, with a ~1.5-fold increased compaction. The observation is consistent with the previous report that distinctly higher compaction in >10 Mb-scale was restricted to specific pairs of X chromosome segments^36^.

We next investigated the geometric features of Xa and Xi. Decode-PAINT enables us to scrutinize the organization of chromosome loci at the nanoscale with high localization precision and sampling efficiency. Our quantitative imaging with Decode-PAINT has revealed that there are no significant differences in geometric parameters between Xa and Xi at the ensemble level (**SFig. 6**; *p* > 0.05, the Mann-Whitney U test, except for the surface-to-volume ratio of S2) or the single-cell pairs (**Fig. 2g–k**; *p* > 0.05, the Wilcoxon signed-rank test).

### Geometric analysis of X chromosome spots at the nanoscale

Finally, we turned our attention to the internal organization of the X chromosomes. While there is no difference in geometric features between Xa and Xi, there could be some differences in the distribution of compactness in the imaged spots. To assess this, we built an analysis method using a voxel-based scanning of the targets (**Fig. 2i and SFig. 7**). We first converted the localization clusters to voxels using a given voxel bin size and then identified voxels exposed to the outer environment (surface voxels) and voxels embedded into the other voxels (core voxels) to calculate the core voxel ratio (**SFig. 7a**). Next, we plotted the core voxel ratio vs. the size of the voxel bin and fitted it with a sigmoid function to obtain the voxel bin size producing the half-maximum occupation of the core voxels (*B*_0_), which represents how the inside of the imaged genomic locus is packed (**Fig. 2i and SFig. 7b**). No significant difference in *B*_0_ was observed between Xa and Xi at the single-cell level (**Fig. 2j**; ~32 nm vs. ~29 nm; p > 0.05, Wilcoxon signed-rank test, n_pairs = 34), suggesting that Xi has a similar internal compaction level with Xa. These observations support the idea that the localized chromosome compaction is not essential for the transcriptional silencing of the Xi. In the future, we anticipate that our approach could be used to investigate other heterochromatic regions in order to assess their local compaction states and the influence of these states on their regulation.

## Discussion

By separating the genomic target identification from single molecule localization, Decode-PAINT allows us to achieve both high throughput imaging of several loci and direct quantitative comparison of super-resolution chromatin structure with a simple and easy-to-implement optical setup. This development opens the door to quantitative super-resolution studies of the nuclear organization at the nanoscale towards in situ single-cell structural genomics. We envision that in combination with the recent advancement in faster DNA-PAINT methods^39–41^ and denaturation-free in situ hybridization^42^, our approach will enable researchers to perform a systems-level analysis of chromatin organization and architecture in individual cells that expect to be broadly applicable to a wide range of biological questions in health and in diseases, such as a comparison between monoallelically expressed genes and conformation changes over heterochromatin formation.

## Materials and Methods

All procedures were performed at room temperature, except where otherwise specified. Step-by-step protocols for probe design, oligonucleotide ordering, PER concatemerization, RNA FISH in cells, and RNA FISH in tissues can be found in the Supplementary Protocols.

### Materials

Lists of oligonucleotide sequences, and library information such as coordinates, barcodes, and density are presented as Supplementary Table 1.

### Buffers

Hybridization buffer: 2× SSC, 10% (wt/vol) dextran sulfate, 0.1% (vol/vol) Tween-20

Wash buffer: 1× PBS, 20% (vol/vol) formamide

Displacement buffer: 1× PBS, 60% formamide

Imaging buffer: 1 × PBS buffer containing 500 mM NaCl, 5× PCA (Sigma-Aldrich, 37580-25G-F), and 2 mM Trolox, 2 U/ml PCD (OYC America, 46852004),

### DNA and RNA FISH probe design

The Oligopaint DNA FISH probe set was designed to target 200-kb genomic spots spanning the entire human X chromosome with 1000 sequences per target. The genomic homology sequences were obtained from the hg38 whole-genome probe sets designed using the OligoMiner pipeline run with the ‘balance’ settings (https://yin.hms.harvard.edu/oligoMiner/list.html), as previously described^30^. Universal forward and reverse-priming sequences for probe preparation and a spot-specific 42-mer sequence for barcoding were appended to each genomic homology sequence. The XIST RNA FISH probe set was designed to target exons of the XIST transcript with 96 sequences. The genomic locations of the exons of the human XIST gene were acquired from the UCSC Genome Browser^43^ and used in combination with the ‘intersectBed’ utility of BedTools^44^ to isolate probe oligonucleotides from the same hg38 whole-genome probe sets above. An imager-binding sequence for detection was appended to the 3’ end of each transcript homology sequence. All Oligopaint FISH probe sequences and coordinates for libraries used in this study can be found in Supplementary Table 1.

### Probe synthesis

The X chromosome DNA FISH probe library was ordered from Twist Bioscience and prepared using T7 in vitro transcription and reverse transcription as previously described^30^. In brief, libraries were first amplified using emulsion PCR, followed by large-scale PCR, and then in vitro transcription to generate RNA complements of probes. Reverse transcription and subsequent digestion of RNA produced single-stranded DNA probe sets. The XIST RNA FISH probe library was ordered from IDT as 96-well plate oligos and mixed.

### Bridge oligo design and preparation

The 42-mer barcode sequences were designed as described previously^30^. In brief, the orthogonal 42-mer sequences were generated computationally and screened using Bowtie, Jellyfish, and BLAST to check the probability of off-target binding to target genomes. NUPACK^45^ was further used to evaluate the single-strandedness and duplexing probabilities of bridge sequences. Two of the DNA-PAINT docking sequences and a PER primer sequence were then appended to the ends of the 42-mer barcode sequences. The bridge oligos were ordered unpurified (standard desalting) from IDT. PER concatemerization of the bridge oligos was performed as described previously^30^. In brief, reactions were prepared in 1X PBS containing 10 mM MgSO4, 0.8 U μl-1 of Bst LF polymerase (NEB, M0275L), 300 μM each of dATP, dCTP, and dTTP (NEB, N0446S), 100 nM of Clean.G hairpin, 50–1000 nM of hairpin and water to 90 μl. After incubation for 15 min at 37 °C, 10 μl of 10 μM primer was added, and the reaction was incubated another 2–3 h, followed by heat treatment at 80 °C for 20 min to inactivate the polymerase. The reactions were purified and concentrated using a MinElute kit (Qiagen, cat. No. 28004) with ultrapure water elution to desalt and reduce volume. See Supplementary Table 2 for the 42-mer bridge sequences and PER hairpins used in this study.

### Cell culture

IMR-90 (human female lung fibroblast; ATCC, CCL-186) cells were grown in DMEM (Gibco, 10564) supplemented with 10% (vol/vol) serum (Gibco, cat. no. 10437), and 1× penicillin-streptomycin (Gibco, cat. no. 15070). Cells were cultured in a 10-cm dish at 37 °C in the presence of 5% CO_2_ and split 1:4 twice weekly. To avoid the accumulation of senescent cells, cells were maintained up to passage 10 before restart.

### Sample preparation for in situ hybridization

All experiments were performed using eight-well chambers (Ibidi, 80827) or channeled chambers (Ibidi, 80168, 0.4 mm height, and 10812). For channeled chambers, a sticky plastic channel and a coverslip were assembled and then incubated at 37 °C overnight before use. IMR-90 cells from ~80% confluent 10-cm dishes were detached from the dishes using 1 ml of trypsin, followed by neutralization with 1.5 ml of fresh medium. The chambers were then seeded with cells at a 1:3–1:4 dilution and incubated at 37 °C in the presence of 5% CO_2_ until ~50% confluency. All the following procedures were performed at 23 °C unless specified. Cells were rinsed twice in 1 × PBS for 1 min, fixed in 1 × PBS containing 4% (wt/vol) paraformaldehyde for 10 min, quenched in 1× PBS containing 100 mM NH4Cl for 5 min, and then rinsed twice in 1× PBS for 1 min. Fixed cells were then permeabilized with 1× PBS containing 0.5% (vol/vol) Triton X-100 for 10 min and then rinsed in 1× PBS containing 0.1% (vol/vol) Tween-20. Samples were immediately subject to FISH to minimize RNA degradation.

### 3D DNA-RNA FISH

After permeabilization, samples were incubated in 0.1 M HCl for 5 min and washed twice in 2× SSCT for 1 min. Samples were then equilibrated in 2×SSCT containing 50% (vol/vol) formamide at 23 °C for 2 min, transferred to fresh 2×SSCT containing 50% (vol/vol) formamide, and incubated at 60 °C for at least 60 min for prehybridization. ISH solution consisted of 2× SSCT containing 50% (vol/vol) formamide, 10% (wt/vol) dextran sulfate, ~800 ng/μl tRNA mix (Sigma-Aldrich, R8508), 200 ng/μl sheared salmon sperm DNA (Invitrogen, AM9680), 800 nM primary X chromosome ISH probes (~47 nM/spot), 800 nM XIST RNA FISH probes, and each PER extended bridge oligo at a final concentration of 80 nM. After denaturation at 78 °C for 3 min on a flatblock thermocycler (Eppendorf, Mastercycler Nexus), samples were incubated at 45 °C for 36–48 hr in a benchtop incubator. Next, 2× SSCT prewarmed at 60 °C was added to samples, and the hybridization solution was aspirated. Samples were then washed four times in 2× SSCT at 60 °C for 5 min, twice in 2× SSCT at 23 °C for 2 min, twice in 1× PBS at 23 °C for 1 min. Gold nanoparticles (40 nm; Sigma-Aldrich, cat. no. 753637) were diluted 1:100 in 1× PBS and deposited on the samples by centrifugation on a swing rotor at 500× g, 23 °C for 3 min to facilitate drift correction. Samples were then washed twice in 1× PBS at 23 °C for 1 min. Samples were optionally stored at 4 °C for 1–2 d before being used.

### Optical setup

SABER pre-decoding and DNA-PAINT imaging were carried out on an inverted custom-built microscope based on a Nikon Ti Eclipse body with the Perfect Focus System, a CFI Apo TIRF 100X Oil (N.A. 1.49) objective, and a piezo Z scanner (PI, P-736) mounted on an XY motorized stage.

For diffraction-limited imaging (XIST RNA FISH and SABER pre-decoding), a Spectra X LED light engine (Lumencor) was used for excitation in a widefield manner with the following settings: (1) a 395/25-nm, 295-mW LED for DAPI signal; (2) a 470/24-nm, 196-mW LED for ATTO 488 signal; (3) a 550/15-nm, 260-mW LED for ATTO 565 signal; and (4) a 640/30-nm, 231-mW LED for Alexa Fluor 647 signal. Emission light was spectrally filtered by one of four filter cubes: (1) Semrock BFPA-Basic-NTE for DAPI; (2) Semrock FITC-2024BNTE-ZERO for ATTO 488; (3) Semrock TRITC-B-NTE-0 for ATTO 565; and (4) Semrock Cy5-4040C-NTEZERO for Alexa Fluor 647 signal. A MicAO 3DSR adaptive optics device (Imagine Optic) was installed downstream of the emission light path to correct optical aberrations from the set-up and the sample.

For DNA-PAINT imaging, a 532-nm laser (MPB Communications Inc., 1 W, DPSS-system) was used for excitation in an oblique illumination manner with a 100 mW input at an effective power density of ~2 kW/cm2. The 532-nm laser beam was passed through a quarter-wave plate (Thorlabs, WPQ05M-532), placed at 45° to the polarization axis, and directed to the objective through an excitation filter (Chroma ZET532/10x) via a long-pass dichroic mirror (Chroma ZT532RDC_UF2). The laser beam diameter was then expanded to ~100 mm using a commercial variable beam expander (Edmund Optics, Broadband VIS 2X-8X) followed by a custom-built Galilean telescope. The laser beam was then coupled into the microscope objective using a motorized mirror to generate an oblique illumination. Emission light was spectrally filtered by a filter cube (Chroma ET542LP and Chroma ET550LP). Emission light was then directed into the MicAO 3DSR adaptive optics device to correct optical aberration and engineer point spread functions (PSFs) for astigmatism-based 3D super-resolution reconstruction^46^. All images were acquired using an Andor Zyla 4.2 Plus sCMOS camera with 6.5-μm camera pixels and rolling shutter readout at a bandwidth of 200 MHz at 16 bit, resulting in an effective pixel size of 65 nm.

### Astigmatism-based 3D reconstruction

To estimate the axial positions of single-molecule emitters, optical astigmatism (−60 nm of “Astigmatism at 0°”) was applied using the adaptive optics device to cause an asymmetric distortion along the Z-axis in the observed PSFs. Image stacks of 100-nm fluorescent beads immobilized on a coverslip surface were acquired in a range of ± 1000 nm with respect to the coverslip with spacing in Z between 5 nm for the calibration of the astigmatism-based axial detection. The image stacks were then processed using Picasso to generate an empirically-derived calibration curve by the least-square polynomial fitting algorithm^47^.

### Exchange-PAINT

For SFig. 1, Exchange-PAINT was carried out by using 1–3 nM of Cy3B-labeled 9-mer imager oligos in the imaging buffer. Before Exchange-PAINT imaging, the Z range of the overall target volume was roughly estimated by detecting ATTO 488 fluorescent signals attached to the 5’ end of the primary FISH probes. For each color, five of 5000 frames with a distance of 170 nm in Z were acquired at a camera exposure time of 200 ms, resulting in a total of 25000 frames in a range of ± ~800 nm with respect to the estimated center of the target volume. Gold nanoparticles (40 nm; Sigma-Aldrich, 753637) were used as fiducial markers to facilitate drift correction. The data were segmented using the DBSCAN clustering algorithm^48^ with input parameters ‘epsilon’ set to 60 nm and ‘minimum number of samples’ to 150 for the data taken with P16 and 100 for the rest of the data. See Supplementary Table 2 for additional details.

### Decode-PAINT

#### XIST RNA imaging and SABER pre-decoding

All the procedures were carried out at 23 °C. Samples were incubated in a fluorescent hybridization solution consisting of 1× PBS and 100 nM fluorescent imager strands for 15 min. Samples were then washed twice in 1× PBS containing 20% formamide for 5 min and rinsed twice with 1× PBS for 1 min. Samples were stained with 1 × PBS containing 1 μg/ml DAPI for 5 min followed by washing twice with 1 × PBS. Detection of the fluorescent signals was carried out in the imaging buffer. Image stacks were acquired in a range from −2 μm to 8 μm with respect to the coverslip surface with a distance of 50–100 nm in Z. After imaging, samples were incubated in the displacement buffer to remove the hybridized fluorescent imager strands. See Supplementary Table 2 for additional details.

#### DNA-PAINT image acquisition

Images were acquired by choosing a region of interest with a size of 800 × 800 pixels (52 × 52 μm). For Fig. 1, DNA-PAINT was carried out by using 1 nM of Cy3B-labeled 9-mer imager oligos in the imaging buffer. Twenty stacks of 5,000 frames with a distance of 170 nm in Z were acquired at a camera exposure time of 150 ms, resulting in a total of 100,000 frames in a range from 200 nm to ~4 μm with respect to the coverslip surface. For Fig. 2, DNA-PAINT was carried out by using 1 nM of Cy3B-labeled 8-mer imager oligos in 1X PBS buffer containing 500 mM NaCl, 2 U/ml PCD (OYC, 46852004), 5X PCA (Sigma-Aldrich, 37580-25G-F), and 2 mM Trolox. Nineteen stacks of 5000 frames with a distance of 85 nm in Z were acquired at a camera exposure time of 100 ms, resulting in a total of 95,000 frames in a range from 200 nm to ~2 μm with respect to the coverslip surface. Prior to acquiring each focal plane, a total of 100 frames of the coverslip surface were acquired to monitor axial stage drift over time. From each single-molecule localization event, ~6,000 photons were collected. Gold nanoparticles (40 nm; Sigma-Aldrich, 753637) were used as fiducial markers to facilitate drift correction. See Supplementary Table 2 for additional details.

#### DNA-PAINT data processing

The lateral and axial positions of single-molecule localization, drift correction, and alignment were carried out using Picasso as described previously^47^. In brief, for each image stack of single-molecule events, the lateral and axial positions of single-molecule localization events were determined using Picasso followed by drift correction. The stacks of single-molecule localization data were aligned using imaged gold nanoparticles as fiducial markers and then merged into a single dataset based on the defined Z-step at imaging. For Fig. 2, the axial position of each focal plane was corrected using the average axial position of gold nanoparticles imaged on the surface prior to merging. The data were segmented using the HDBSCAN clustering algorithm^49^ with the input parameter ‘minimum cluster size’ set to 30 and 40 for Figs. 1 and 2, respectively. Global XY and XZ localization precisions were estimated by the nearest neighbor analysis (NeNA) using Picasso. FRC resolution analysis for Fig. S4 was carried out using Vutara SRX (Bruker) based on localizations without linking.

#### Image processing for diffraction-limited images

For diffraction-limited images (XIST RNA and SABER), image processing was carried out using ImageJ/Fiji^50^. The Z stacks of XIST RNA and SABER images were first aligned to the cognate DNA-PAINT reconstructed images based on the fiducial markers. Next, for each channel, the maximum intensity value of each pixel along the Z-axis was projected onto the 2D plane. The maximum intensity projections were then cropped to the selected ROI (~256 × 256 pixels), covering all the SABER signals. Yen’s thresholding^51^ was applied to the maximum intensity projections to convert them to pre-decoding binary filters.

#### Decoding single-molecule localization clusters

The decoding was carried out using custom Python scripts (**SFig. 3a**). *Overlap score O(α,β)* of an HDBSCAN cluster *α* and a pre-decoding binary filter of the genomic spot *β* is defined as follows (**SFig. 3b**):

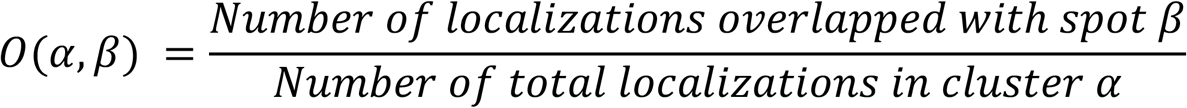

First, overlap scores were computed for all pairs of HDBSCAN clusters and pre-decoding binary filters (9 for Fig. 1 or 6 for Fig. 2). Given an HDBSCAN cluster *x,* if only the pre-decoding binary filter of the genomic spot *y* gives the maximum of *O(x, y)* and the score is equal to or greater than 0.5, then *x* is assigned to spot *y*. If there is no *y* that gives *O(x, y)* equal or greater than 0.5 (e.g., not overlapped), then x is rejected. If there is more than one filter that gives the maximum *overlap score* (e.g., a cluster fully overlapped with more than one filter gives a tie), then x is set to “unassigned.”

### Data analysis

#### Volume, surface area, and dimensionless surface-area-to-volume ratio

For Fig. 2h and i, the point cloud data of localizations was converted into 3D binary voxels with 50-nm cubic binning. Volume was calculated by counting the number of total voxels containing at least one localization and multiplying it by the volume of a voxel. The surface area was calculated by counting the number of total open faces of the voxels and multiplying it by the area of a face. For Fig. 2j, a dimensionless version of surface-area-to-volume (SA:V^2/3^) was calculated using surface area and volume^2/3^.

#### Gyration radius

The gyration radius (*R_g_*) of the imaged chromatin segment is defined as:

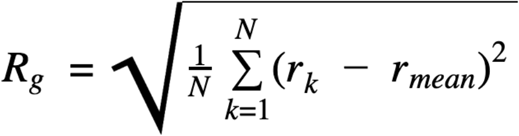

where *N* is the number of total localizations, *r_k_* is a vector of the XYZ coordinate of an individual localization, and *r_mean_* is the mean position of all *N* localizations.

#### Analysis of internal compaction level using voxel scanning

For Fig. 2i, j, and SFig. 7, only single-cell paired spots that have 1000 or more localizations were analyzed (n_spotpair = 34). The point cloud data of localizations of each Decode-PAINT imaged spot was converted into 3D binary voxels with 10–65-nm cubic binning in 5-nm increments. Next, each voxel was classified based on the number of its faces in contact with neighboring voxels into two categories: (i) surface or (ii) core (**SFig. 7a**). *Core voxel ratio* is defined as the number of surface voxels divided by the number of total voxels. Obviously, the core voxel ratio approaches 0, where the voxel bin size goes too small (e.g., 10 nm), as each localization is converted to an isolated surface voxel. In contrast, the core voxel ratio saturates, where the voxel bin size goes too large (e.g., 65 nm). The core voxel ratio was plotted against the voxel bin size, and the plot was fitted by a sigmoid function as:

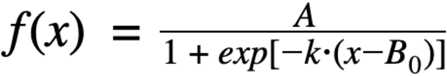

where *A* is the maximum value of the function, *k* is the slope of the function, and *B_0_* is the voxel bin size giving the half maximum of the function, which represents the apparent compaction level of the point cloud data. It should be noted that the core voxel ratio declines again, where the voxel bin size goes over the Nyquist frequency of the data. Therefore, the fitting was performed in the range of voxel bin size between 10 nm and the voxel bin size giving the maximum of the core voxel ratio within the range of the plot (10–65 nm).

## Data availability

All data are available in the main text or the supplementary materials. Materials and unprocessed image data are available upon request. Information regarding all datasets can be found in Supplementary Table 2.

## Code availability

All code is available at https://github.com/beliveau-lab/Decode-PAINT

## Acknowledgments

This work was supported by the National Institutes of Health (under grants 1R01EB018659-01 to P.Y., 1UG3HL145600 to P.Y., 1R01GM124401 to P.Y., 1U01MH106011-01 to P.Y., and 1DP1GM133052 to P.Y.), the Office of Naval Research (under grants N00014-16-1-2410 to P.Y. and N00014-18-1-2549 to P.Y.), the National Science Foundation (under grant CCF-1317291 to P.Y. and a Graduate Research Fellowship to J.Y.K.), the Damon Runyon Cancer Research Foundation (under a fellowship to B.J.B.), the Uehara Memorial Foundation (under a fellowship to H.M.S.), and the Wyss Institute’s Molecular Robotics Initiative (MRI) (P.Y., H.M.S., J.Y.K., and B.J.B.).

## Author contributions

H.M.S., B.J.B., and P.Y. conceived the study. H.M.S., J.Y.K., and B.J.B. designed oligo probes and bridge oligos. H.M.S. designed and executed imaging experiments, developed the analysis pipeline and methods, and analyzed data. J.Y.K. and C.-t.W. contributed to optimizing and performing data analysis. H.M.S., C.-t.W., B.J.B., and P.Y. wrote the manuscript. All authors edited and approved the manuscript. B.J.B. and P.Y. supervised the work.

---

**Figure S1.**
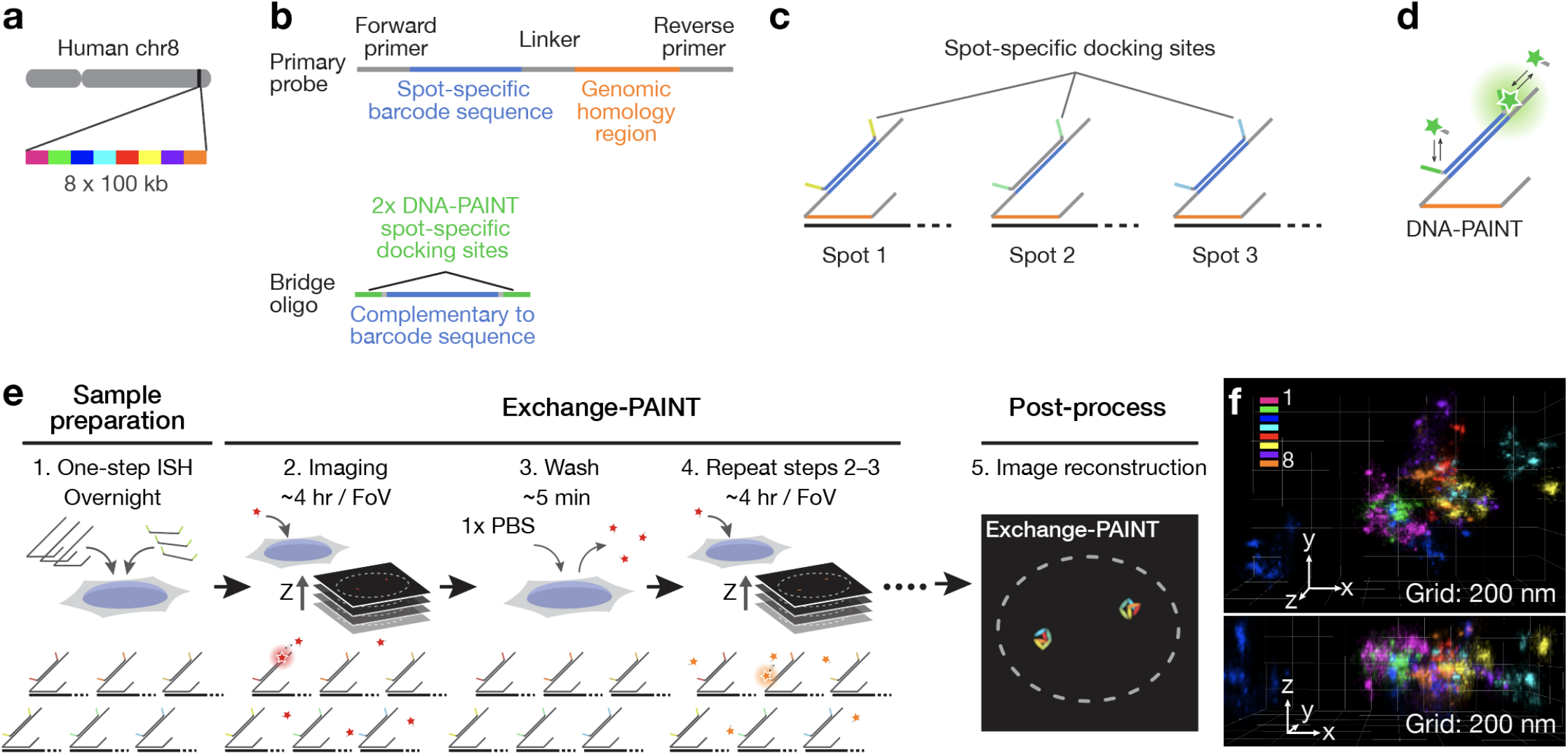
Exchange-PAINT with FISH. **a**, Schematic of eight targeted regions along human chromosome 8. Each set of probes per spot had orthogonal 42-nucleotide oligomer barcode sequences appended to their 5’ ends. Eight 42-nucleotide oligomer bridge sequences with two identical copies of eight orthogonal DNA-PAINT docking sequences were simultaneously hybridized. The genomic length of each target is 100 kb. **b**, Primary probe and secondary bridge oligo designs for Exchange-PAINT FISH. The primary probe design is the same as **Figure 1a**. A secondary bridge oligo carries a complementary sequence to a spot-specific barcode and two copies of a spot-specific DNA-PAINT docking sequence. **c**, The tiling strategy for exchange-PAINT with FISH. A unique DNA-PAINT docking sequence is assigned to each genomic target spot. **d**, DNA-PAINT readout from secondary bridge oligos. The bridge oligos generate single-molecule signals via transient binding between DNA-PAINT imagers and docking sites. **e**, Schematic overview of Exchange-PAINT FISH. First, primary probe oligos and bridge oligos are hybridized to genomic targets in fixed samples following standard ISH protocols. After washing, each targeted spot is imaged with sequential DNA-PAINT using orthogonal imagers. DNA-PAINT data are aligned using fiducial markers and then converted to a reconstructed image. **f**, A representative image of 8-color Exchange-PAINT FISH on human chromosome 8.

**Figure S2.**
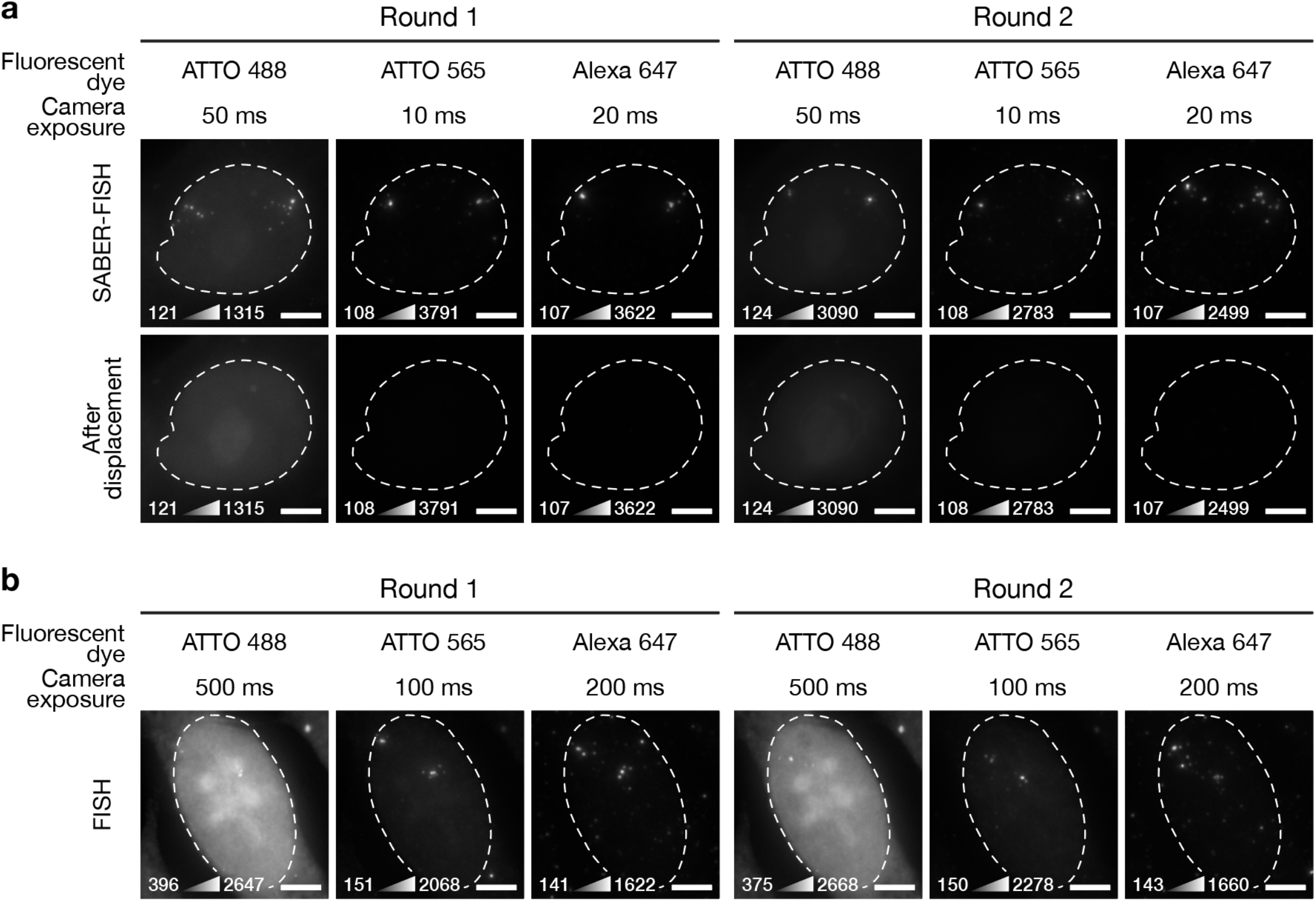
SABER pre-decoding validation. **a**, Two rounds of Exchange-SABER for pre-decoding. Short camera exposure time (≤ 50 ms) is enough to achieve a high signal-to-noise ratio (SNR). After the displacement of SABER imagers, no residual signals were observed. **b**, Two rounds of DNA exchange imaging with unextended bridge oligos. The experiment was carried out with the same procedure for Figure 2a but with unextended bridge oligos. While SNR was comparable to SABER images for ATTO 565 and Alexa 647, 10-fold longer camera exposure time was needed. For ATTO 488, SNR was worse than SABER images due to the high nucleus background. For each image, the minimum and maximum pixel intensity value used to set the display scale is indicated in the lower left. Dashed line, nucleus. Scale bar, 5 μm.

**Figure S3.**
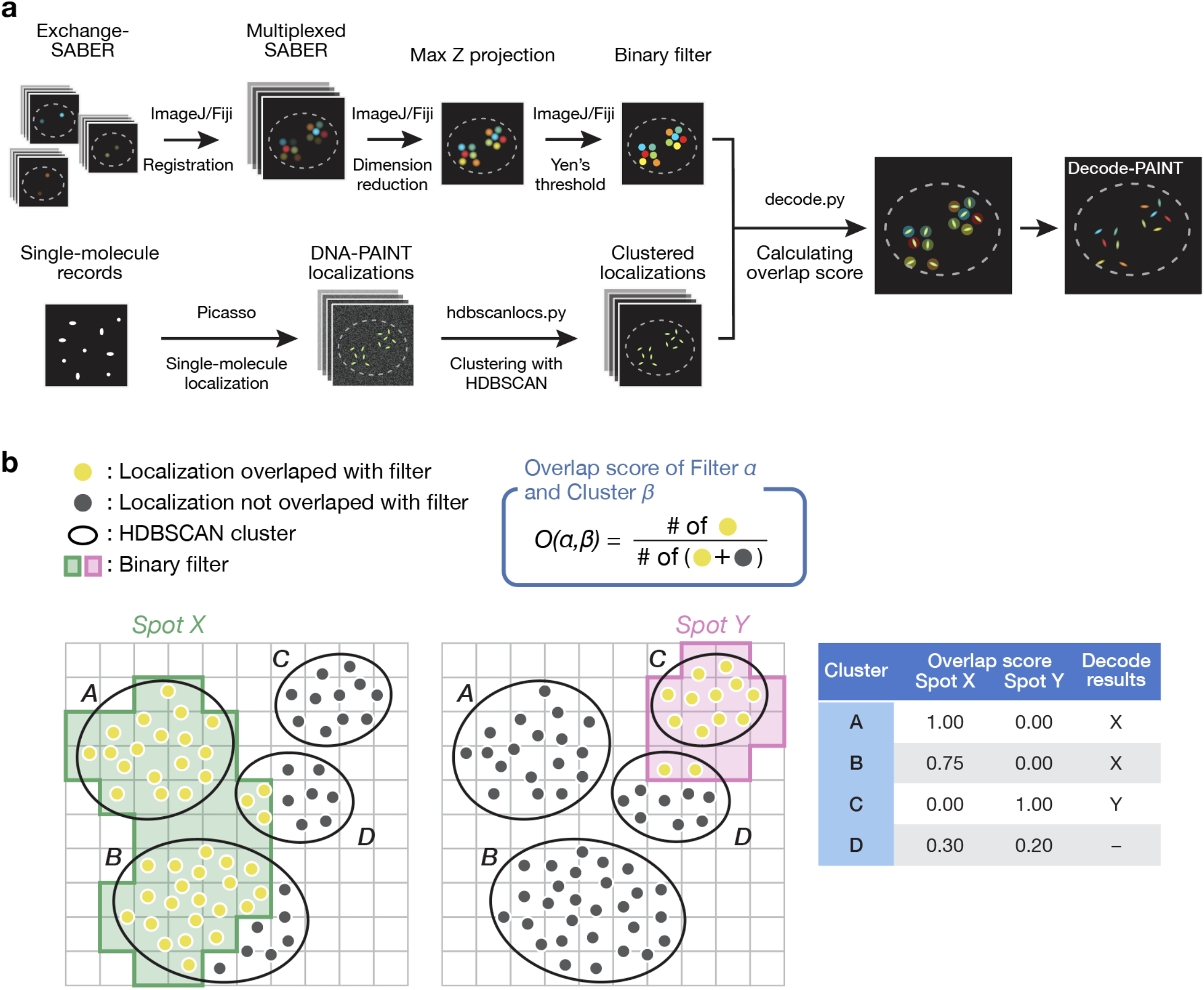
Implementation of decoding. **a,** Schematic overview of the decoding workflow. **b**, Schematic overview of overlap score computation. See *Decoding single-molecule localization clusters* for the detail.

**Figure S4.**
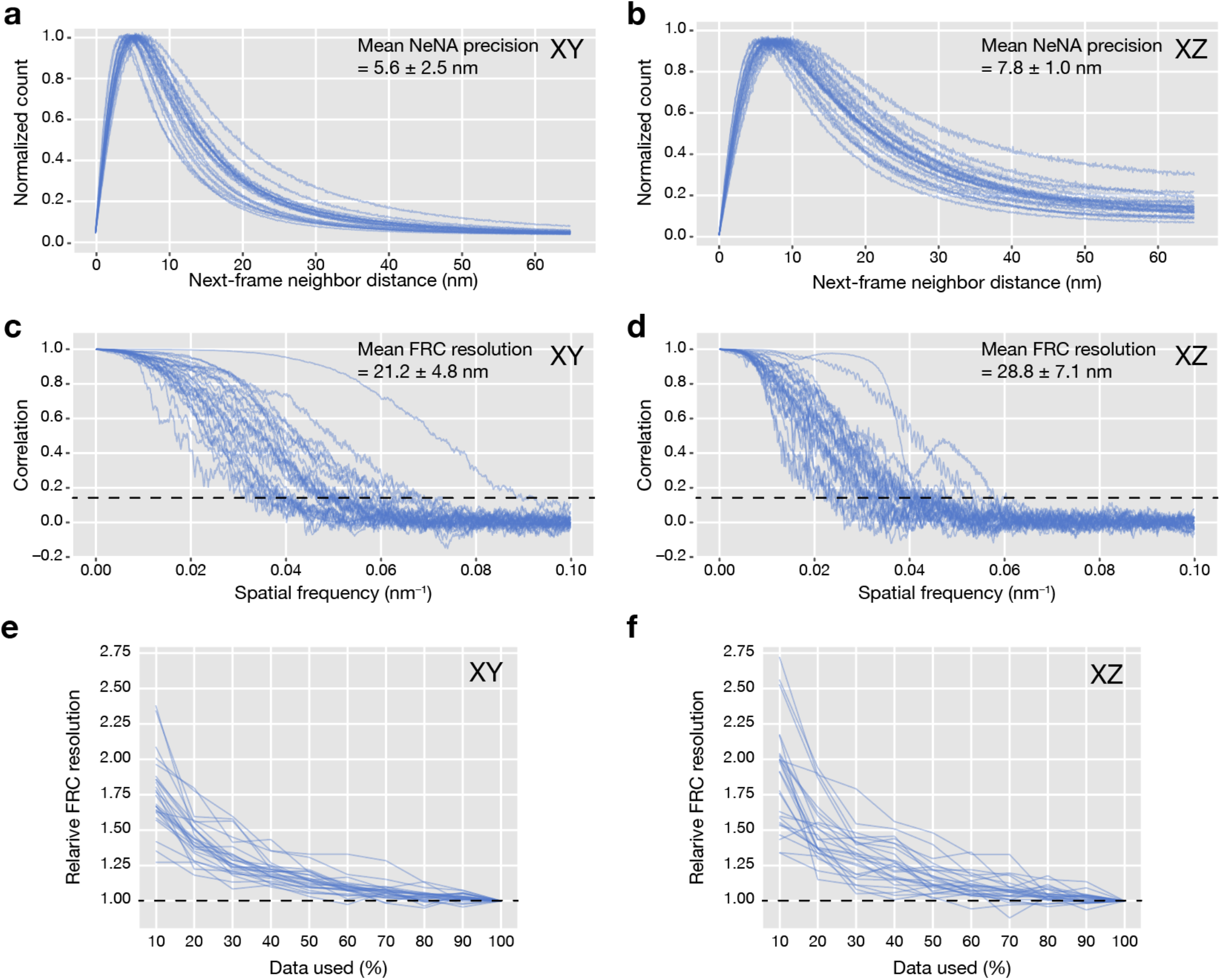
Nearest neighbor-based analysis (NeNA) for localization precision and Fourier ring correlation (FRC) analysis for structural resolution. **a,b**, NeNA localization precision plots for XY (**a**) and XZ (**b**) generated from the datasets for Figure 2. Mean NeNA localization precision ± s.d. for XY and XZ is 5.6 ± 2.5 and 7.8 ± 1.0 nm, respectively. **c,d,** Fourier ring correlation resolution analysis for XY (**c**) and XZ (**d**) generated from the datasets for Figure 2. Mean FRC resolution with 0.143 (1/7) cutoff ± s.d. for XY and XZ is 21.2 ± 4.8 and 28.8 ± 7.1 nm, respectively. **e,f**, Saturation plot of relative FRC resolution calculated using downsampled datasets to assess data sampling efficiency.

**Figure S5.**
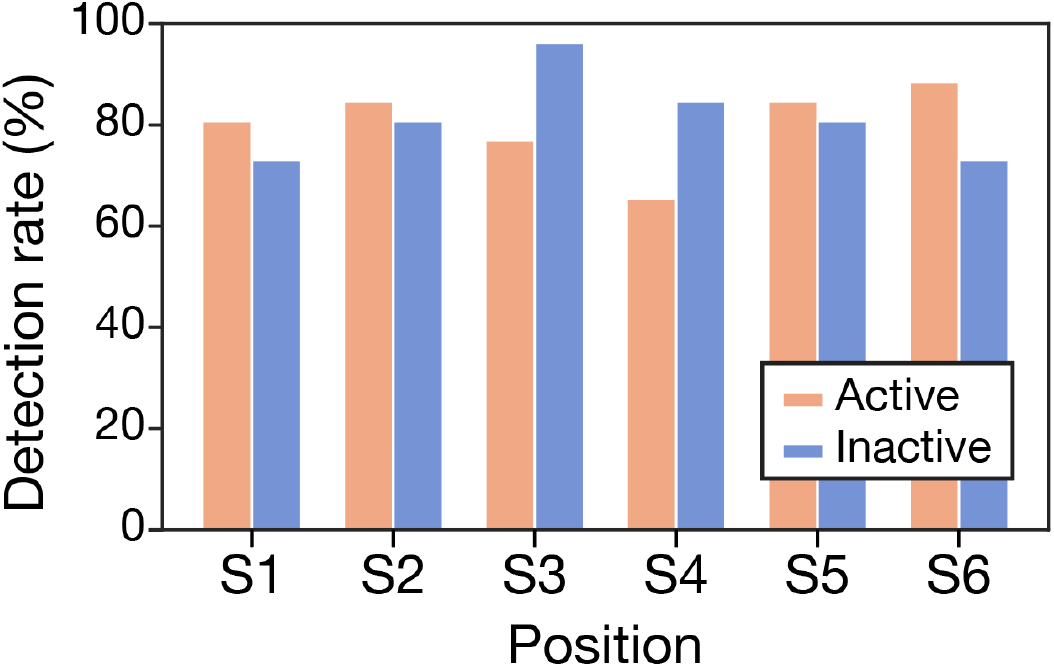
Detection rate of 6-color Decode-PAINT on the X chromosome. Targeted regions were detected in ~60–90% of the imaged cells (n = 26) from 3 replicates. Some regions were not detected mostly because those were out of the Z range covered by our microscopy modality.

**Figure S6.**
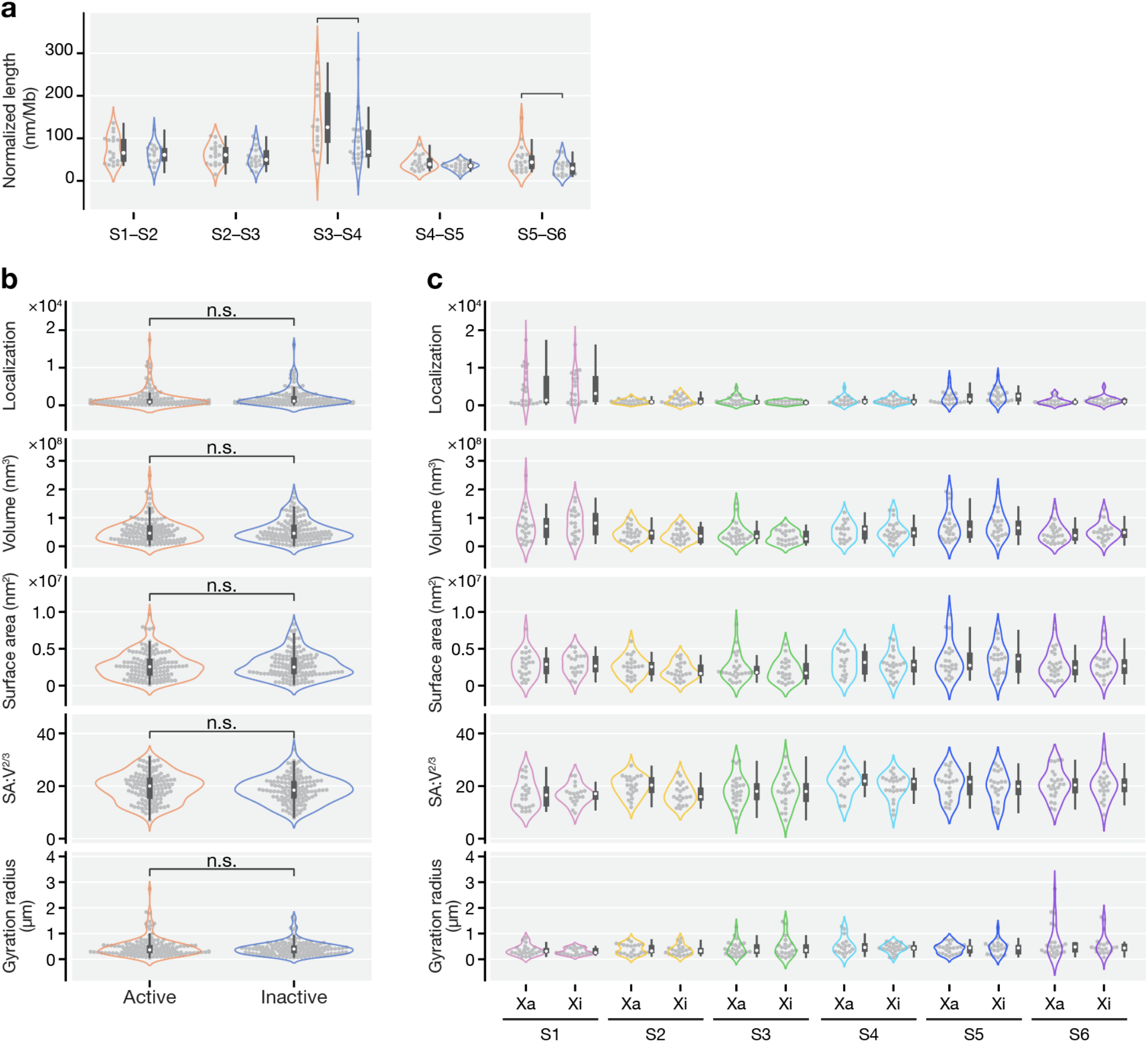
Unpaired comparison of the geometric features between active and inactive X chromosomes. **a**, Violin plots of normalized 3D distances of adjacent spot pairs between active and inactive X chromosomes. The data plotted is the same as in Fig. 2f, however, single-cell pairs are disregarded. Swarms of observations are overlaid on top of the violin plots. The miniature box plot in or next to each violin plot shows the interquartile range (IQR), the white dot marks the median, and the thin lines extend to show 1.5 × IQR. **b,c**, Violin plots of geometric features between active and inactive X chromosomes without (**b**) or with (**c**) consideration of genomic positions. The data plotted is the same as in Fig. 2g–k, however, single-cell pairs are disregarded.

**Figure S7.**
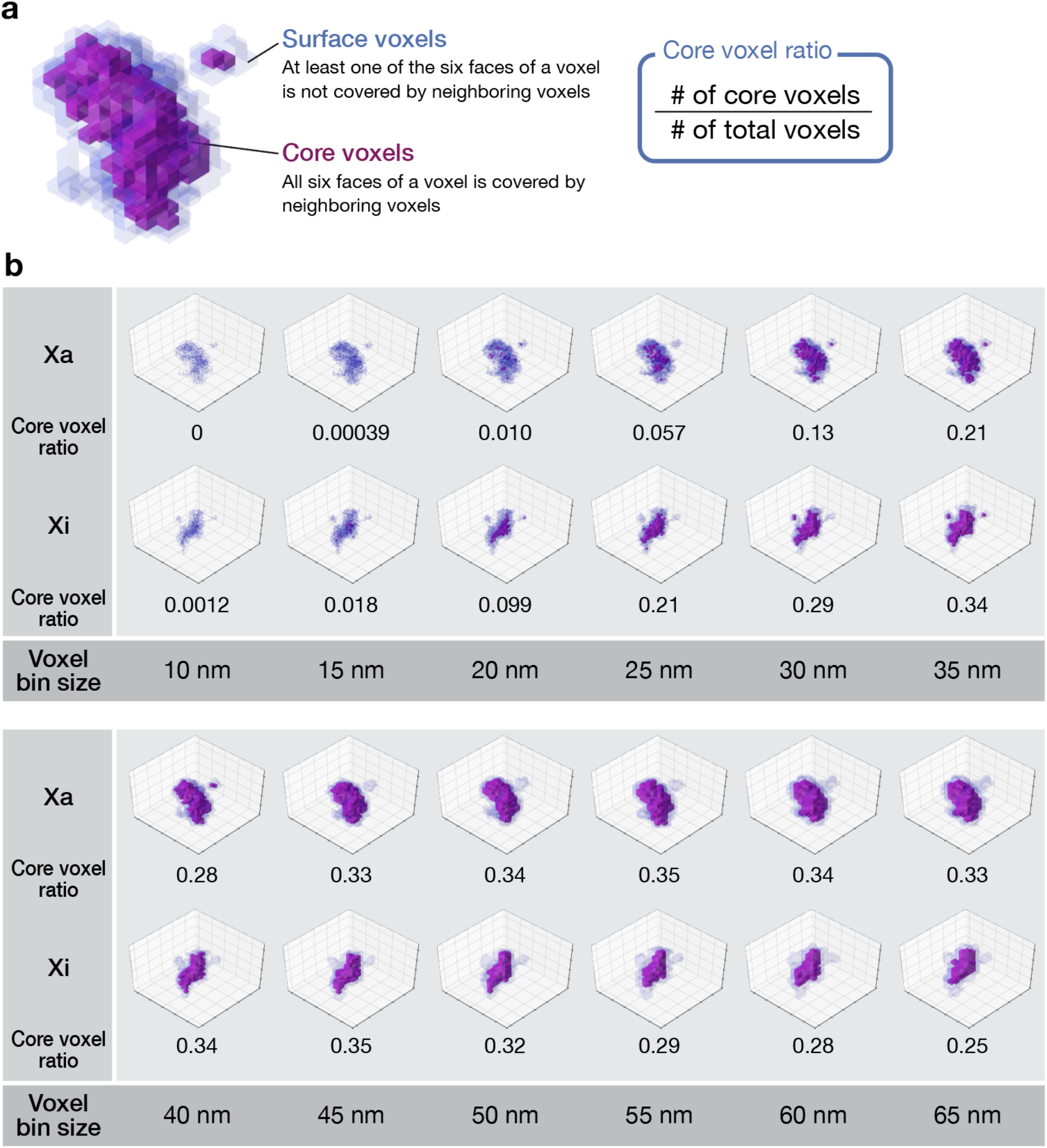
Implementation of voxel-scanning analysis. **a**, Schematic of the core voxel ratio. **b**, Voxel-scanning analysis. A representative data of S5 on the active and inactive X chromosomes in a single cell was shown.

